# Sex-related differences in vascular remodeling of sodium overloaded normotensive mice

**DOI:** 10.1101/2024.07.09.602349

**Authors:** Katia A. S. Viegas, Juliane C. S. Silva, Rariane S. Lima, Cintia T. Lima, Natalia N. Peron, Maikon B. Silva, Maria Claudia Irigoyen, Silvia Lacchini

**Affiliations:** Department of Anatomy, Institute of Biomedical Sciences, University of Sao Paulo (USP), Sao Paulo, Brazil; Hypertension Unit, Heart Institute, University of Sao Paulo Medical School (FMUSP), Sao Paulo, Brazil

**Keywords:** Sodium overload, gender, vascular collagen deposition, arterial stiffening

## Abstract

*Background and Aims:* Primary hypertension affect about 20% of adults in developed societies and, associated with high salt intake, leads to vascular remodeling, an adaptive physiological response of blood vessels that driven by repair, inflammation, or cell growth, is but may contribute to vascular diseases over time. This study examined vascular remodeling in the aorta and cardiac arteries, focused on gender-specific responses to sodium overload. *Methods and Results:* Adult male and female C57Bl/6 mice were divided into six groups: control with filtered water (Cont M; Cont F), 1% NaCl for two weeks (Salt-2 M; Salt-2 F), and 1% NaCl for twelve weeks (Salt-12 M; Salt-12 F). Blood pressure (BP) and heart rate (HR) were measured using tail plethysmography, and metabolic cages recorded 24-hour water intake and urine output. Morphometric analysis of the aorta and cardiac arteries included assessments of elastic laminae and collagen fibers, using Weigert van Gieson and Picrosirius staining. No changes in BP and HR were observed. Sodium intake increased water consumption in both genders after two weeks, but only males showed increased urine output. Vascular responses differed: males exhibited delayed increases in aortic elastic lamellae, while females showed earlier changes. Elastic lamellae in cardiac arteries remained unchanged. Collagen deposition increased in aortic walls for both genders but decreased by 50% in male cardiac arteries. Males showed thick collagen fibers, while females had thin and thick fibers. *Conclusion:* High sodium intake caused arterial stiffness through distinct mechanisms in males and females, even in normotensive animals.

## 1. INTRODUCTION

Cardiovascular diseases are the main cause of morbidity and mortality in the world. According to the World Health Organization (WHO), 17.3 million people died in 2008 from diseases related to heart and blood vessels which represents two-thirds of all deaths worldwide. In addition, around 16.5% of deaths that affect the cardiovascular system have been associated with arterial hypertension, including 51% from stroke and 45% from coronary heart disease deaths [1]. Primary hypertension is a multifactorial condition arising from the complex interplay between genetic predisposition and environmental factors, including stress, sedentary lifestyle, obesity, and excessive dietary salt intake. It is among the most prevalent cardiovascular diseases, affecting approximately 20% of the adult population in developed countries [2]. Left unmanaged, primary hypertension can accelerate the atherosclerotic process, increasing the risk of coronary artery disease [3,4].

In 2019, the average daily salt consumption among adults aged 25 and older in the region was estimated at 8.5 grams per day, with men consuming approximately 9.5g and women 7.1g. These values surpass the body’s physiological needs and are 1.7 times greater than the WHO’s recommended limit of less than 5g salt/day (equivalent to 2g sodium/day) [1,5]. The increasing salt intake is detrimental to health and presents different consequences depending on the population heterogeneity and experimental design [6]. Recent data on sodium intake show that populations around the world are consuming much more sodium than is physiologically necessary, there are regional variations in sodium intake [7,8]. The global average sodium intake is estimated at 3.95g/day, based on a meta-analysis of research from 187 countries using 24-hour urine collections [9].

Variations in the vascular microenvironment or arterial injuries, which can be caused, for example, by excess sodium, can induce changes in the structure and function of the cardiovascular system, triggering a complex and dynamic process described as vascular remodeling that aims to meet the physiological needs of local blood supply as occurs, for example, in the mesenteric arteries [10–12]. with the endothelium also playing a crucial role in vascular homeostasis [13–15].

Research with rats subjected to sodium overload for 2 and 12 weeks demonstrated that it modifies perivascular collagen deposition, increasing thick fibers and decreasing thin fibers [16]. The vascular remodeling emerges from situations of repair, inflammation, development or cell growth [17,18]. The extracellular matrix (ECM) is a three-dimensional network formed by numerous proteins such as proteoglycans, glycosaminoglycans, elastin, laminin and matrix metalloproteases although collagen has been considered the main component [19–22]. The ECM is responsible for maintaining the tissue integrity in the vessel besides regulating the cell migration, and also acting as a reservoir of cytokines and growth factors [21,23–25].

The tissue perfusion is promoted by microcirculation where vessels, especially arteries, are capable to change the diameter, structure, and composition of the vascular wall in order to increase the surface area needed for blood exchange. This response is essential for tissue integrity and organ function. The perfusion of tissue varying on the metabolic requirements and depending on the stimulus may lead to remodeling for the purpose of maintain the stability between blood flow and shear stress [26,27].

Vascular remodeling is a physiological adaptation that may lead to vascular diseases over time. Studies suggest women are more prone to cardiovascular diseases than when, though this difference diminishes is postmenopausal women [28–31]. Additionally, women exhibit distinct cardiac and vascular remodeling compared to men, prompting ongoing research into the cardioprotective effect of female hormones, which remain incompletely understood [12,32,33].

Considering that cardiovascular diseases are the leading cause of morbidity and mortality, with rising hypertension rates and increased dietary sodium consumption worldwide, this study hypothesizes that dietary sodium may affect vascular system adaptability, regardless of blood pressure, with responses varying by exposure duration and gender. It aims to analyze vascular remodeling in large and small arteries (aorta and cardiac arteries) and compare male and female mice under sodium overload.

## 2. MATERIAL AND METHODS

### 2.1 Animals

The present study used adult male and female C57Bl/6 mice, obtained from the animal care unit of the Department of Anatomy of the Institute of Biomedical Sciences at the University of Sao Paulo. The animals received standard laboratory chow and water *ad libitum* and were distributed in cages in a temperature-controlled room with a 12-h dark-light cycle. The animals were randomly assigned to one of six groups: control, receiving filtered water (Cont M; Cont F); 1% NaCl solution to drink for two weeks (Salt-2 M; Salt-2 F); 1% NaCl solution to drink for twelve weeks (Salt-12 M; Salt-12 F), according to the previous study [16]. The treatments started on the 4-week-old mice and were continued until they reached 16 weeks of age although the salt-2 group received 1% NaCl solution in the drinking water in the last 2 weeks of protocol, therefore, all animals were of the same age at the end of treatment (Figure 1). Experiments were conducted according to the ethical principles established by the Ethics Committee on the Use of Animals of the Institute of Biomedical Sciences at the University of Sao Paulo, research protocol 049/2011.

**Figure 1.**
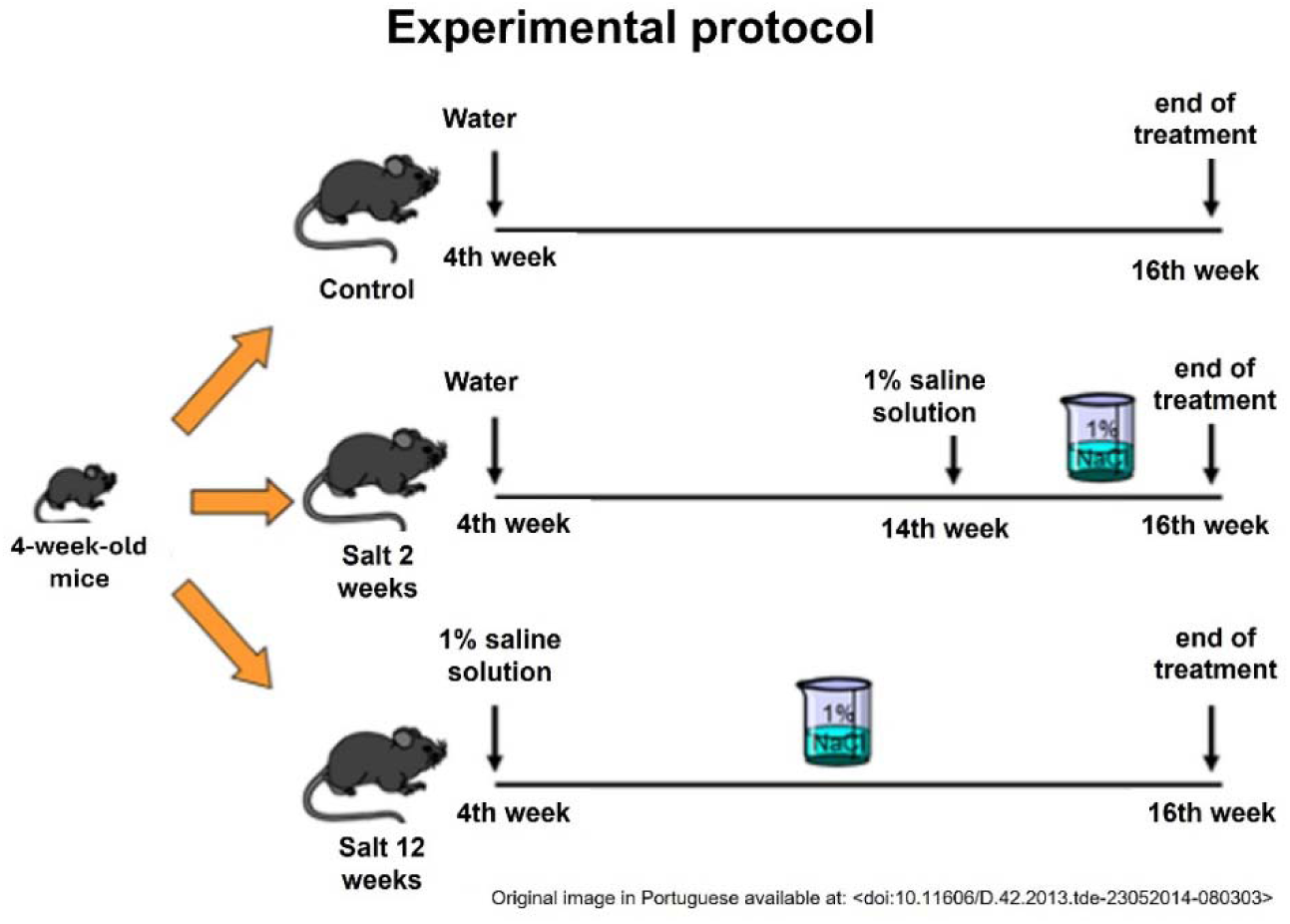
Experimental protocol diagram

### 2.2 Blood Pressure and Heart Rate assessment

In the last week of treatment, pulse blood pressure (BP) was measured by tail plethysmography. The animals were adapted to the system during the week prior to the measurement. BP and heart rate (HR) were measured 10 times for two consecutive days. Peak systolic BP was captured by the BP-2000 series II system, according to the previous study [16].

### 2.3 Water intake and urine output measurements

On the last week (2^nd^ and 12^th^ week) of treatment, mice were individually housed in metabolic cages for a single period of 24 hours and provided with tap water (Cont group) or 1% NaCl water (Salt-2; Salt-12 groups) and regular chow *ad libitum*. During one day prior to the measurement, the animals were adapted to the system. Water intake and urine output were measured within the 24 hours in the metabolic cage.

### 2.4 Histological Evaluation

#### 2.4.1 Tissue collection and preparation

At the end of treatment, the animals were euthanized with an anesthetic overdose of ketamine (180 mg/kg) supplemented with xylazine (20 mg/kg) and were subsequently perfused with 0.9% NaCl solution followed by a buffered 4% formalin solution. After that, the tissues were harvested and maintained in formalin during a period of 24h. Following the fixation process, tissues were processed and paraplast embedded for histological evaluation.

#### 2.4.2 Morphometry

Elastic laminae and collagen fibers in the aorta and cardiac arteries, stained with Weigert van Gieson and Picrosirius, respectively, were analyzed to assess extracellular matrix components responsible for vascular compliance and resistance. Blinded morphometric analyses of 5-µm tissue sections were conducted using a Zeiss Axio Scope II microscope with image analysis software. Intima and media areas were calculated based on differences in elastic lamina boundaries, and collagen fiber deposition was quantified under polarized light, comparing mature, intermediary, and immature fibers.

### 2.5 Statistical Analysis

All values are expressed as means±SD. These mean values of each animal were compared by Analysis of Variance (ANOVA) of two ways (gender and time of treatment), complemented by the Bonferroni’s test. For all statistical analyses, p≤0.05 were considered statistically significant. The program GraphPad Prism, version 5.01 (GraphPad Software, Inc., 2007) was used to perform the statistical analysis.

## 3. RESULTS

### 3.1 Blood Pressure, Heart Rate, Water Intake and Urine Output measurements

The treatment with 1% NaCl solution did not promote any change in the hemodynamic parameters evaluated. There was no difference in values for systolic arterial pressure and heart rate in sodium overloaded mice compared to the control group (Figure 2A and B).

**Figure 2.**
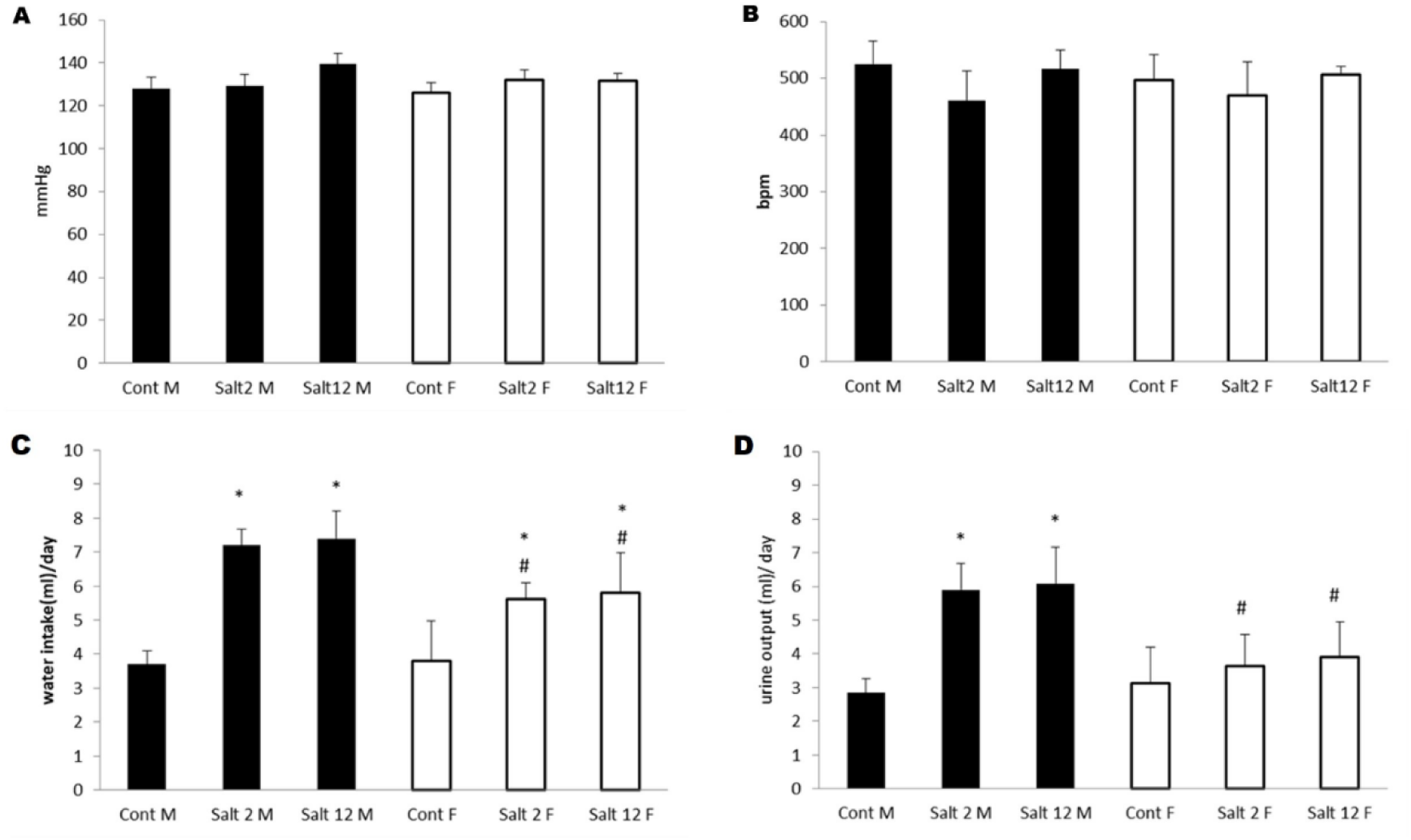
Tail cuff plethysmography showing mean values systolic pressure (A) and heart rate (B), with no differences between genders or treatments. (C, D) Analysis for water intake and urine output in 1% NaCl-treated groups: increased water intake (C) after two weeks in both sexes and increased urine output (D) in males. male (M) and female (F); N=10/group; *p≤0.05 vs. control; #p≤0.05 vs. M.

Treated mice showed an increase in water consumption, with intake rising in both sexes after two weeks and stabilizing after 12 weeks (Figure 2C and 2D). Male mice also had higher urine output, unlike females, who showed no significant change. Female exhibited a smaller increase than males in both metrics across the treatment period.

### 3.3 Elastic lamellae analysis

Morphometric analysis revealed that 1% NaCl in drinking water altered aortic structure in both sexes. Male showed increased elastic lamellae after 12 weeks, while females responded earlier, with lamellae increasing after 2 weeks and nearly doubling by 12 weeks of treatment (Figure 3).

**Figure 3.**
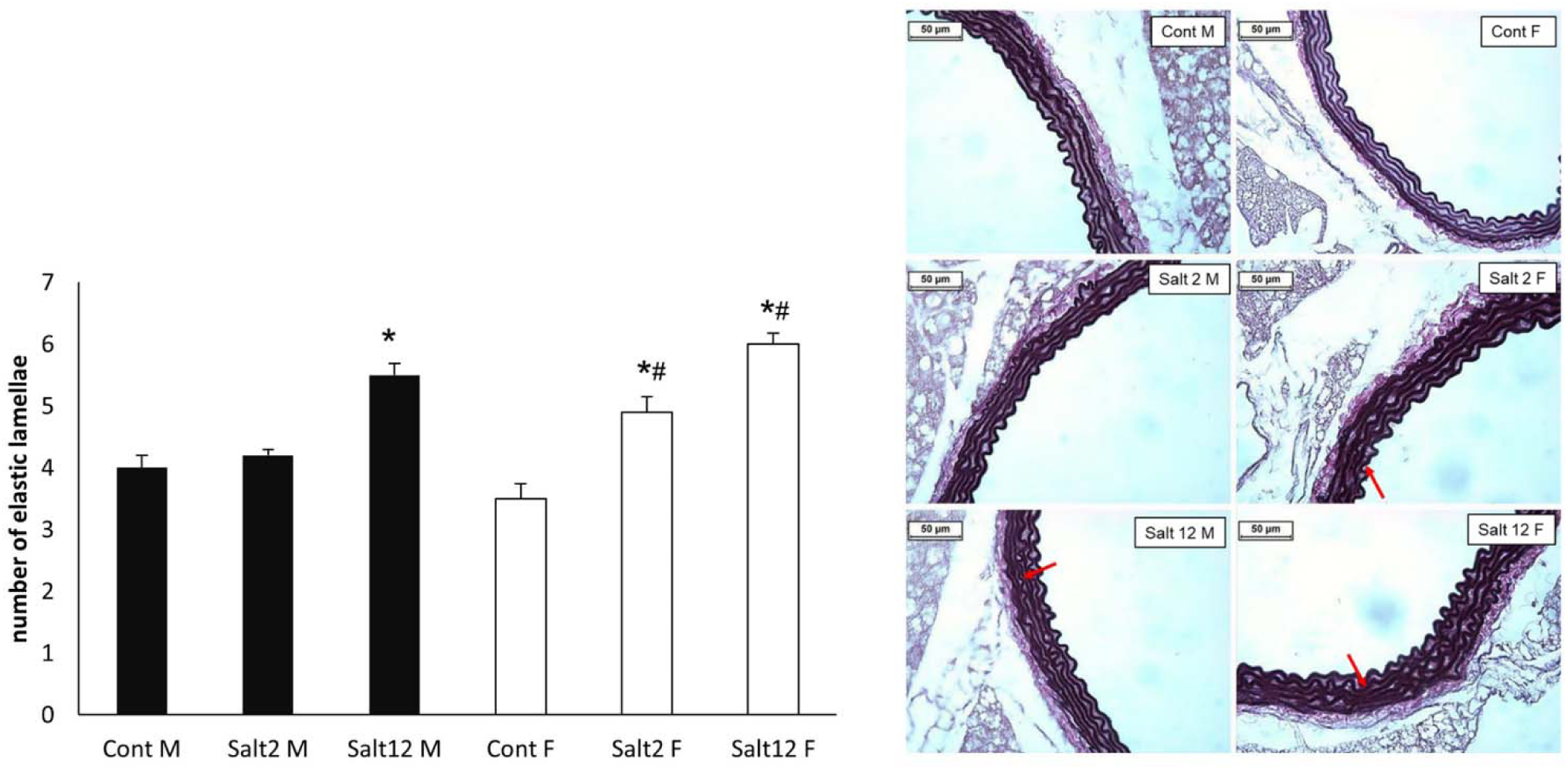
Number of elastic lamellae in aorta, showing higher increase in the number of elastic lamellae (red arrows). Saline overload increased the number of elastic lamellae in male (M) after chronic treatment, while female (F) increased also after shorter time. N=10/group; * p≤0.05 compared to control group; # p≤0.05 compared to respective male group. Staining: Weigert van Gieson.

Unlike the aorta, cardiac arteries showed no change in elastic lamellae regardless of treatment duration or sex. All groups had only the inner elastic laminae, consistent with expectations, as these arteries regulate vascular resistance by altering smooth muscle cell content or ECM production, particularly collagen fibers.

### 3.4 Collagen fibers deposition

To assess treatment effects on vessel structure and vascular resistance, we analyzed perivascular collagen deposition. In the aorta, collagen increased gradually in males (+30.2% at 2 weeks, +36% at 12 weeks) and in females (+54.5% at 12 weeks), with males showing a more pronounced early response. However, females had higher collagen levels after chronic sodium intake (+51.4%), indicating greater vascular stiffness (Figure 4A).

**Figure 4.**
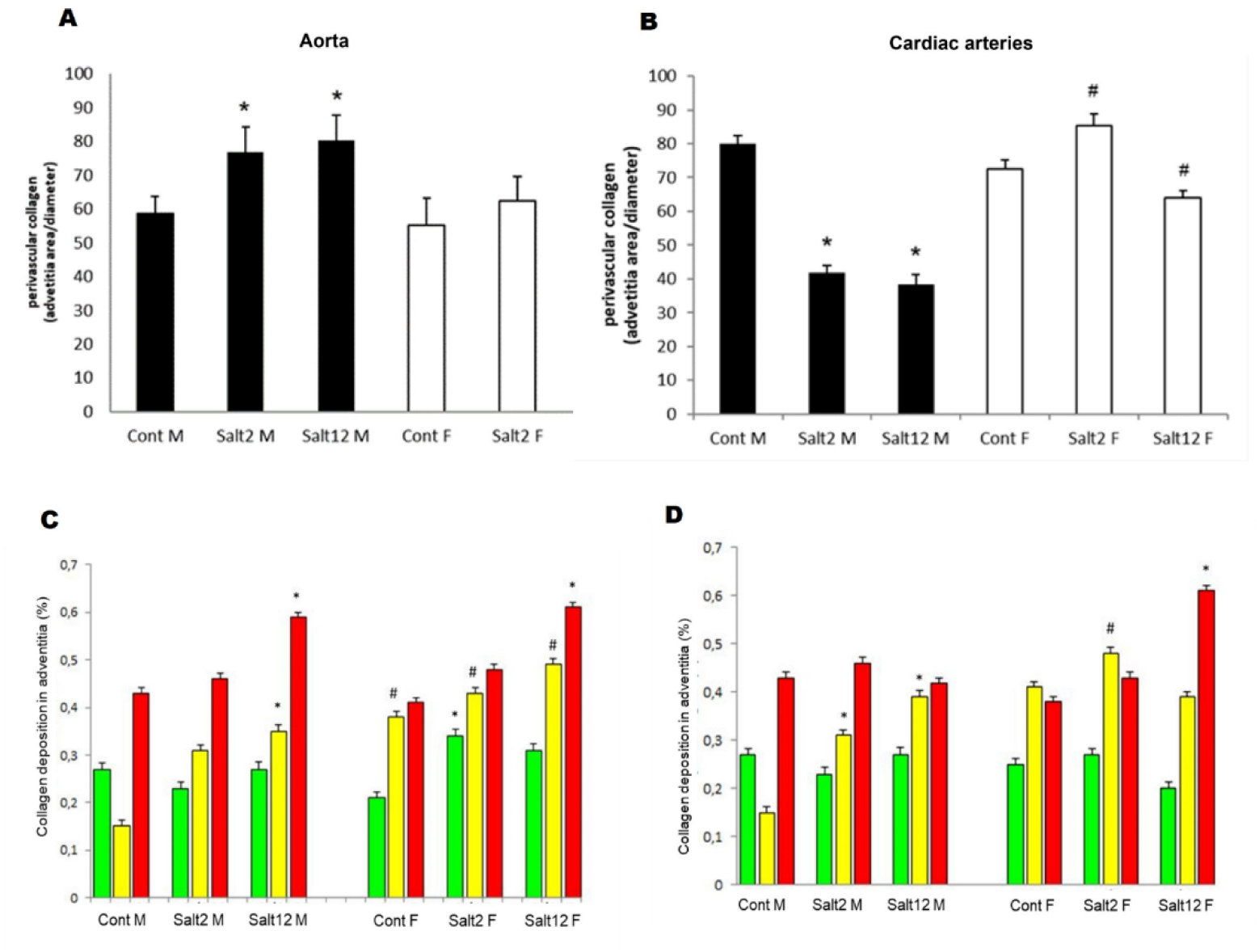
**A-B:** Perivascular collagen deposition in aorta and cardiac arteries. Collagen increased in male (M) aorta with saline, while females (F) rose only after chronic treatment. Cardiac arteries: males decreased collagen; females unchanged. **C-D:** Collagen fibers types in aorta and cardiac arteries: thin (green), intermediary (yellow), thick (red). Aorta: more intermediary/thick fibers (saline) in M; increased with both treatments in F. Cardiac arteries: reduced response in M; increased intermediary (Salt2) and thick fibers (Salt12) in F. *N = 10/group;* **\*** *p* ≤ *0.05 vs. control;* **#** *p* ≤ *0.05 vs. males*.

On the other hand, analyzing collagen fibers deposition in cardiac arteries, we found differences regarding the treatment as well as gender. Males reduced their collagen fiber content 47.8% after 2 weeks of treatment, and even more after 12 weeks (−52.1%). Interestingly, females seemed to have a fluctuation in collagen fiber deposition, presenting a small increase after 2 weeks (+17.4%) and a small reduction (− 11.7%) at the end of treatment; however, these changes were not significant (Figure 4B).

Collagen fibers deposition was also studied under polarized light since it is possible to identify collagen fibers by their refringence. As it can be seen in figure 6, the number of intermediary collagen fibers doubled in the aorta of the male group treated for 12 weeks. Likewise, there was also an increase in the deposition of mature (thick) collagen fibers (Figure 4C). The female group showed a significant increase in the content of intermediary collagen fibers in the aorta when compared to the male group at the same time of the treatment, as well as in young (thin) and mature (thick) collagen fibers after 2 and 12 weeks of treatment, respectively, when compared to the control (Figure 4D).

**Figure 5.**
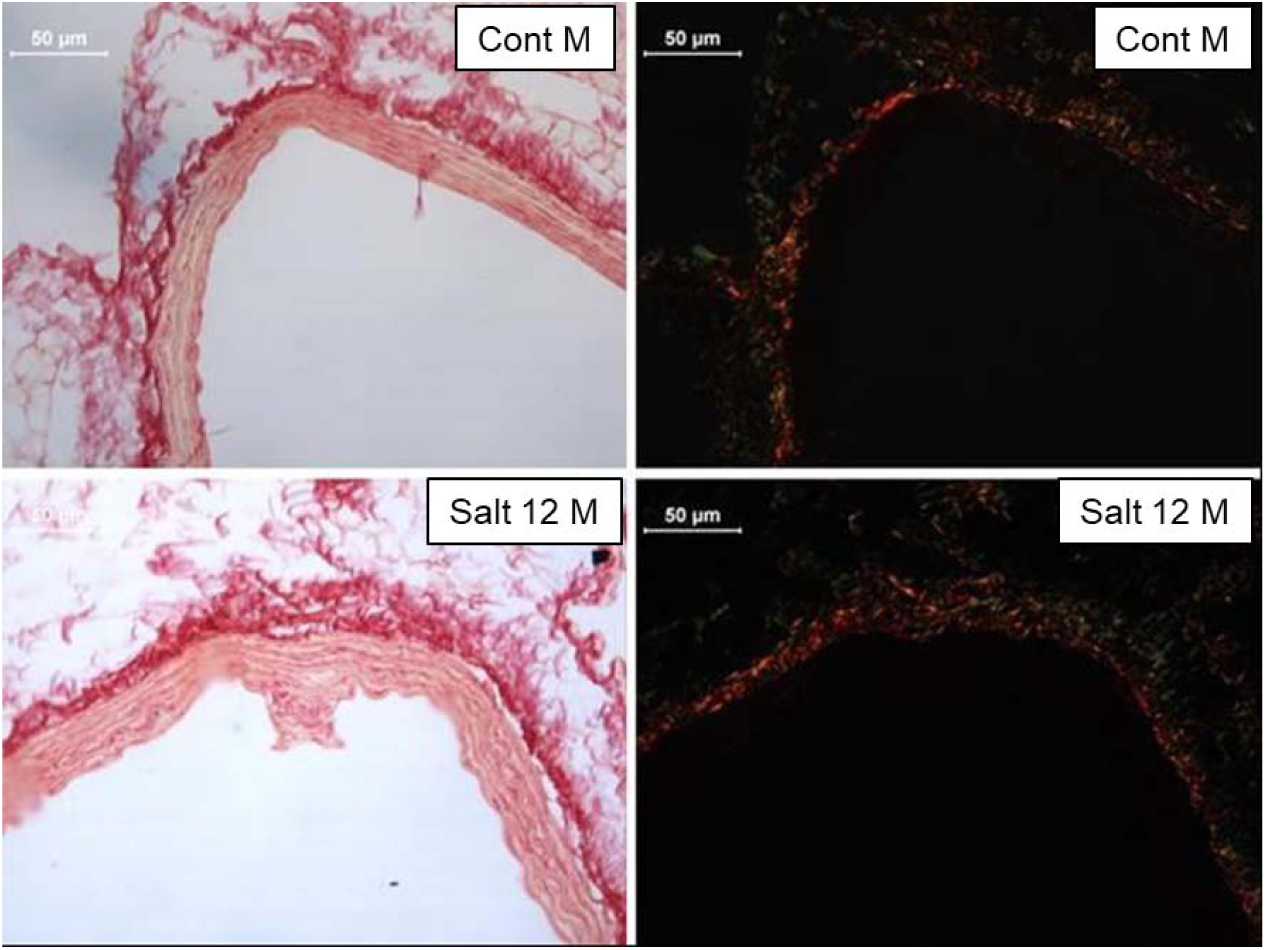
Representative photomicrographs of collagen deposition in the aorta, comparing male (M) and female (F) mice after 12 weeks of high sodium intake. Picrosirius red-stained sections assessed by bright field (left column) and polarized light (right column) in the aorta.

**Figure 6.**
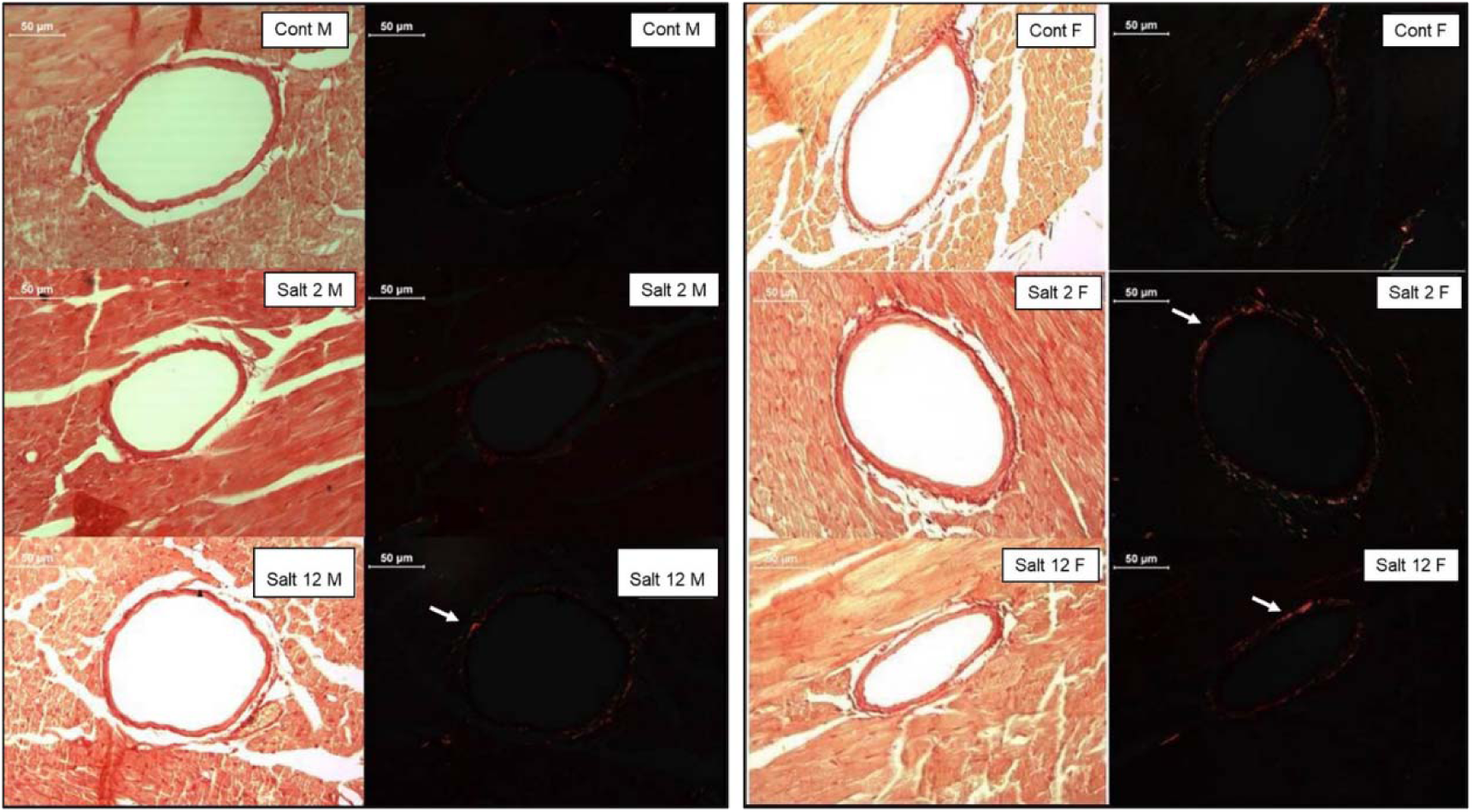
Collagen deposition cardiac arteries analyzed by Sirius red-stained. Sections under bright field (left) and polarized light (right). Polarized light revealed increased thick collagen fibers (white arrow) after 12 weeks of saline treatment in male (M) and showed increased thick collagen fibers in both saline groups in female (F).

Polarizes light analysis of cardiac arteries showed that while perivascular collagen decreased in males, intermediary fibers (yellow) increased significantly with sodium intake (Figures 4C-D, 5, 6A-D). In females, mature (thick) fibers increased later (12 weeks), but intermediary fibers rose earlier (2 weeks) and more prominently than in males (Figures 4 and 6).

## 4. Discussion

During the last decades, several studies have related sodium intake with the onset of cardiovascular diseases. The higher the sodium amount in the diet is, the higher the probability to develop cardiovascular diseases is the opposite has also been confirmed [34–36]. Researches have also found an increase in the mortality index in rats treated with high salt diet [37] and increased blood pressure and endothelial dysfunction [38,39]. Paradoxically, the relationship between sodium intake and cardiovascular disease in some studies has not necessarily been associated with increased blood pressure [40,41].

This study aimed to determine if increased sodium intake induces vascular changes related to arterial pressure and gender. A 1% NaCl diet was chosen based on previous studies in our lab, which showed that even a small salt increase stimulates vascular factor production [16,41,42]. Moreover, higher levels of salt intake will promote gastrointestinal problems that influence the body adaptation, and also the expression of inflammatory markers [43–45]. In this pattern, the age of the animals was also chosen based on studies that showed that the age would influence salt resistance: the older the animal was, the higher the propensity to be salt-sensitive was [46,47]. At the end of treatment, the animals reached 4 months of age; therefore, they were still considered young.

Hence, as described in previous studies, we did not observe any change in blood pressure [16]. However, fewer studies have evaluated the effect of sodium intake in mice, although it is well known that the intake of NaCl 1% in the drinking water did not promote changes in blood pressure of rats after 1 [41] or 6 weeks [42], as well as after 12 months of treatment [16].

The results obtained in the metabolic cage showed that the increase in sodium consumption promoted a significant increase (corresponding to 95% and 100%) in the water intake in males after 2 and 12 weeks, respectively. On the other hand, the females presented a smaller increase of 47% and 53% for the same period. These results are according to other studies that have shown an increase in water intake in animals treated with high salt diet for a short period and also chronically [48].

The animals also increased the urine output; however, only in the male group, an increase of 107% and 114% was observed after 2 and 12 weeks of treatment, respectively. On the other hand, females had a slight increase of 21% and 30% in the volume of urine for the same period. A smaller increase in water intake in females when compared to males may be related to metabolic differences promoted by sexual hormones [49–51]. Although both males and females had increased water intake as well as urine volume excretion, males increased the volume retention by 51% and 53% after 2 and 12 weeks, respectively, while females had a greater increase in the volume retention of 145% and 138% after 2 and 12 weeks, respectively.

The treatment with 1% NaCl also promoted important vascular changes in both males and females, even though it depended on the time of exposure to the diet. These results are according to previous studies which have confirmed that the high salt diet can promote vascular and renal alterations in normotensive or hypertensive subjects [52,53].

The observed changes occurred in elastic and resistance vessels, for example, the aorta, well known for being an elastic artery, increased its elasticity in males treated with 1% NaCl after 12 weeks. Nevertheless, in females, this increase is more precocious at just 2 weeks of treatment. Moreover, analyzing the perivascular collagen fibers in the aorta, we found that males tend to have an increase in these fibers, especially mature (thick) collagen fibers which may promote stiffness as shown in previous studies[40]. Thus, the aorta seems to increase its elasticity in order to counterbalance the higher stiffness. A similar response is shown in females that present not only a greater deposition of mature, but also intermediary collagen fibers suggesting even more stiffness in this vessel.

Resistance vessels, such as cardiac arteries, respond differently from elastic vessels. No change in elastic laminae was observed, as expected for resistance vessels. In males, collagen fiber deposition decreased, while intermediary fibers increased, possibly compensating to stabilize the vascular wall, a pattern shown in previous carotid artery studies [54]. In contrast, females increased deposition of mature and intermediary fibers during treatment, leading to greater stiffness.

In fact, the literature describes sex differences in appetite for salt, where women exhibit a greater preference, especially during pregnancy [50,51,55,56]. Additionally, evidence indicates that sex chromosomes and hormones influence blood pressure regulation, cardiovascular risk factors, and comorbidities differently in men and women [57,58]. Our results suggested a heightened predisposition and poorer vascular response in women under stress factors, such as a high-salt diet.

## 5. CONCLUSION

The results of this study suggest that a 1% increase in NaCl in drinking water of mice did not significantly change blood pressure levels after 2 or 12 weeks of treatment. Therefore, the results did not indicate the hypertension onset. However, the diet may have induced different adaptive vascular responses in males (elastic arteries) and females (resistance arteries), leading to arterial stiffness even in normotensive animals.

Therefore, elucidating the mechanisms underlying these sexual differences in vascular response to salt intake may contribute to the development of gender-specific prevention and therapeutic strategies for cardiovascular diseases.

## Authors contributions

SILVA, J.C.S., VIEGAS, K.A.S., LACCHINI, S. contributed to conceptualization, methodology, writing-review and editing; LIMA, R.S., LIMA, C.T., PERON, N.N., SILVA, M.B. contributed to formal analysis and investigation; IRIGOYEN, M.C. contributed to supervision, LACCHINI, S. contributed to supervision and project administration. All authors contributed to the article and approved the submitted version.

## Declaration of generative AI and AI-assisted technologies in the writing process

During the preparation of this work the authors used ChatGPT in order to reduce characters number in a few paragraphs. After using this tool, the authors reviewed and edited the content as needed and takes full responsibility for the content of the publication.

